# FBL17 targets CDT1a for degradation in early S-phase to prevent Arabidopsis genome instability

**DOI:** 10.1101/774109

**Authors:** Bénédicte Desvoyes, Sandra Noir, Kinda Masoud, María Isabel López, Pascal Genschik, Crisanto Gutierrez

## Abstract

Maintenance of genome integrity depends on controlling the availability of DNA replication initiation proteins, e.g., CDT1, a component of the pre-replication complexes that regulates chromatin licensing for replication. To understand the evolutionary history of CDT1 regulation, we have identified the mechanisms involved in CDT1 dynamics. During cell cycle, CDT1a starts to be loaded early after mitotic exit and maintains high levels until the G1/S transition. Soon after the S-phase onset, CDT1a is rapidly degraded in a proteasome-dependent manner. Plant cells use a specific SCF-mediated pathway that relies on the FBL17 F-box protein for CDT1a degradation, which is independent of CUL4a-containing complexes. A similar oscillatory pattern occurs in endoreplicating cells, where CDT1a is loaded just after finishing the S-phase. CDT1a is necessary to maintain genome stability, an ancient strategy although unique proteins and mechanisms have evolved in different eukaryotic lineages to ensure its degradation during S-phase.

**Impact statement:** The DNA replication protein CDT1a is crucial for genome integrity and is targeted for proteasome degradation just after S-phase initiation by FBL17 in proliferating and endoreplicating cells of Arabidopsis

## Introduction

Faithful genome replication is a complex and risky process that requires robust and highly-regulated overlapping mechanisms to prevent genome instability. They operate primarily at the stage of DNA replication initiation at two complementary levels: selection and activation of DNA replication origins (ORIs), which is known as licensing, and dynamics of initiation proteins.

ORI selection depends on the stepwise assembly of pre-replication complexes (pre-RC) at genomic sites that can potentially be used as active ORIs. The mechanism of sequential assembly of pre-RC components as well as the genomic sites preferred for pre-RC binding are in general terms highly conserved in all eukaryotes. Genomic locations of ORIs possess locally enriched stretches of GC and GC skew, in most of the cases also embedded in an open chromatin landscape typical of proximal promoters, transcriptional start sites and the 5’-end of genes (Cadoret et al., 2008; Costas et al., 2011; Pourkarimi et al., 2016; Rodriguez-Martinez et al., 2017; Sequeira-Mendes et al., 2019). Pre-RCs are macromolecular entities consisting of the origin recognition complex (ORC) heterohexamer that binds DNA and nucleates pre-RC assembly, the (cell division cycle 6) CDC6 and (Cdc10-dependent transcript 1) CDT1 proteins, and the minichromosome maintenance (MCM) heterohexamer (Riera et al., 2017; Yeeles et al., 2015).

Contrary to the relatively high conservation of the general ORI features, regulation of the availability and dynamics of pre-RC protein components is highly species-specific, since multiple mechanisms have been reported in different model systems. Although Orc1 and Cdc6 protein levels vary during the cell cycle (DePamphilis, 2005), Cdt1 is redundantly regulated at multiple levels, revealing its relevance to maintain genome stability and prevent re-replication-associated defects. The first regulatory level corresponds to a transcriptional control using the Cdc10 transcription factor in *Saccharomyces pombe* (Whitehall et al., 1999) whereas in animal and plant cells it relies on the E2F pathway (Castellano et al., 2004; Desvoyes et al., 2006; Yoshida and Inoue, 2004). In addition, Cdt1 availability is regulated during the cell cycle after the G1/S transition. While in *S. cerevisiae* Cdt1 is exported out of the nucleus (Tanaka and Diffley, 2002), in mammalian cells it is targeted to the proteasome for degradation by two E3 ubiquitin ligases that work independently, SCF^Skp2^ and CRL^Cdt2^ (Havens and Walter, 2011; Nishitani et al., 2006). It is noteworthy that targeting by SCF^Skp2^ requires CDK-mediated phosphorylation whereas CRL^Cdt2^ instead requires interaction with PCNA (Havens and Walter, 2011; Li et al., 2003). A third regulatory layer relies on the inhibition of Cdt1 function by geminin (Maiorano et al., 2000; Whittaker et al., 2000; Wohlschlegel et al., 2000), a protein present at high levels in S-phase until it is degraded by the anaphase promoting complex/cyclosome (APC/C) in mitosis (McGarry and Kirschner, 1998).

Maintaining correct levels of Cdt1 is crucial to prevent genome instability both in animals (reviewed in (Blow and Gillespie, 2008)) and plants (Castellano et al., 2004; Raynaud et al., 2005). Given the different mechanisms evolved to control Cdt1 levels and their species-specificity we wondered whether CDT1 function is regulated by ancient mechanisms evolved in plants, as suggested by early observations (Castellano et al., 2004). Here, we have addressed the question of how CDT1 protein levels are controlled in plant cells by analyzing the cell cycle dynamics of Arabidopsis CDT1a and by identifying the features shared with animal cells, such as its fast degradation after the G1/S transition and its targeting for degradation by the FBL17 F-box protein. We have also compared the CDT1a availability in proliferating and endoreplicating cells. The use of live imaging and new marker lines have been instrumental to obtain results that contribute to a better understanding of the evolutionary history of CDT1 regulation, a crucial licensing step for the maintenance of genome stability.

## Results

### Expression of CDT1a in proliferating cells

The Arabidopsis genome contains two genes, *CDT1a* (At2g31270) and *CDT1b* (At3g54710), which encode CDT1 proteins, a key component of the pre-replication complexes (pre-RCs). Transcriptional GUS reporter lines revealed that both *CDT1a* and *CDT1b* genes show a very similar pattern of preferential expression in proliferating tissues (Castellano et al., 2004). Here we have focused on CDT1a, which is an essential protein as revealed by the lethal phenotype of the corresponding T-DNA insertion mutant (Domenichini et al., 2012).

To study the expression dynamics of CDT1a at the protein level we generated a translational fusion of the genomic *CDT1a* locus fused to CFP under its own promoter. Based on our analysis of the chromatin configuration at the CDT1a genomic locus (Sequeira-Mendes et al., 2014), we cloned a promoter region of 1823 bp, encompassing proximal promoter region (state 1 and 2) and the intergenic region chromatin state 4 (Figure 1–figure supplement 1). Constitutive CDT1a overexpression leads to altered levels of nuclear ploidy (Castellano et al., 2004). Thus, we assessed that the *pCDT1a*::*CDT1a-CFP* line, both in a wild type and in the *cdt1a*-*1* mutant backgrounds, showed a nuclear ploidy profile similar to that of the wild type plants, with only a small increased frequency of higher ploidy nuclei (Figure 1A). In fact, the lethal phenotype of *cdt1a-1* is suppressed by expression of the CDT1a-CFP fusion protein (Figure 1–figure supplement 2). These data led us to conclude that the CDT1a-CFP reporter could be used to study the CDT1a dynamics and regulation in proliferating cells of a developing organ.

**Figure 1.**
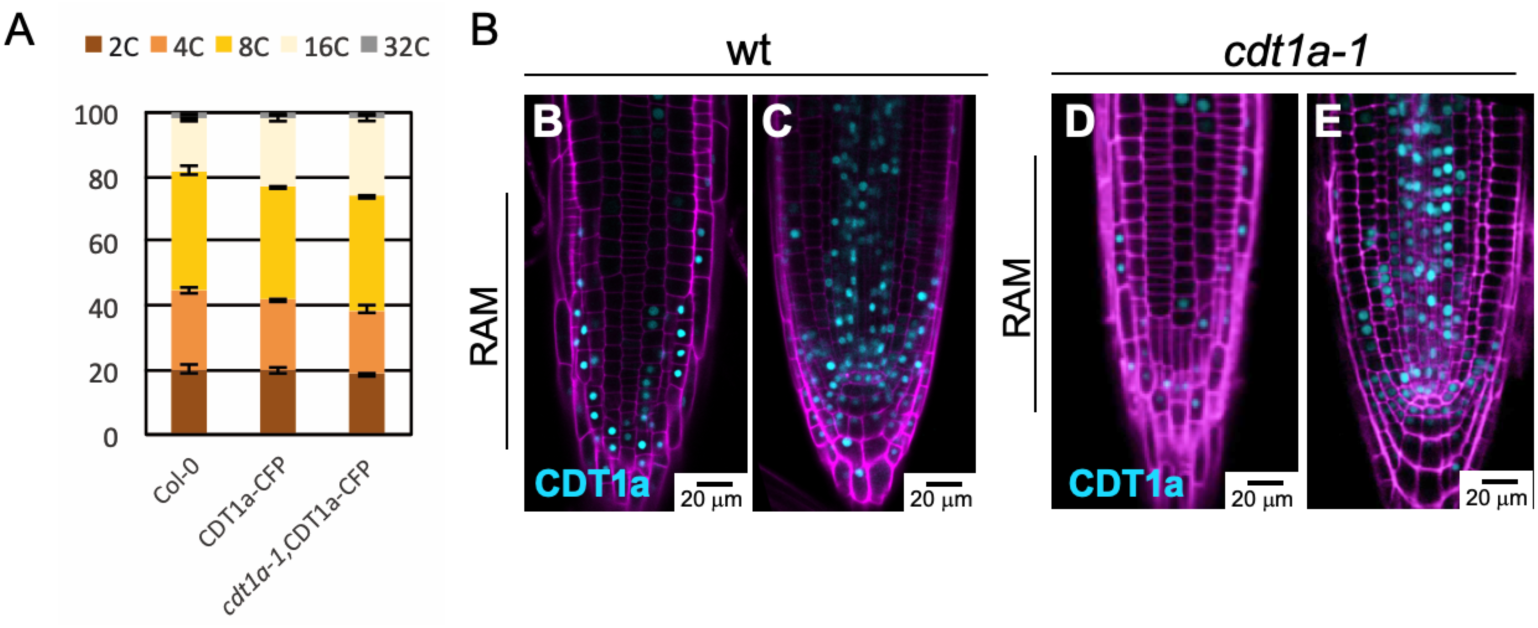
Expression of CDT1a in proliferating cells. (A) Nuclear DNA ploidy profiles of leaves #1/2 of 3-week-old Arabidopsis seedlings. N=10,000 nuclei in each nuclei were scored. Data correspond to mean ± s.d. Differences between ploidy levels of each genotype were not statistically significant using a t test with a Welch correction. (B-E) Detection by confocal microscopy of CDT1a-CFP protein in the root meristem of 5-day-old Arabidopsis seedlings in the wild type (B,C) *cdt1a-1* mutant (lethal in homozygosis; D,E) backgrounds, in the epidermal cell layer (B,D) and in the plane of the quiescent center (C,E).

Analysis by confocal microscopy showed that CDT1a is detected in the different proliferating cells of the root apical meristem (RAM) as a nuclear protein (Figure 1B,C). It is noteworthy that expression of CDT1a-CFP in the *cdt1a-1* mutant background showed an expression pattern similar to that of CDT1a-CFP in the wild type (Figure 1D,E). Epidermal cells showed a clear patchy pattern, typical of cell cycle regulated proteins (Figure 1B,D). Cells of the quiescent center (QC) appear in most cases devoid of CDT1a-CFP signal whereas most of the surrounding stem cells contained CDT1a-CFP (Figure 1C,E). CDT1a-CFP-containing cells were frequent in the stele (Figure 1C,E).

### CDT1a protein is loaded in early G1 and degraded at the G1/S transition in proliferating cells

We used live imaging to study CDT1a dynamics during the cell cycle in the growing Arabidopsis root meristem cells by following the CDT1a-CFP signal in plants also expressing the constitutively expressed HTR5 histone H3.3-mRFP (Otero et al., 2016). Since mitotic cells did not contain CDT1a-CFP we started to follow the progression of mitotic cells into the next cell cycle. We observed that detectable levels of CDT1a appeared ∼25-30min after finishing cytokinesis and continued to accumulate for a few hours during G1 of daughter cells (Figure 2A–movie supplement 1), showing a high degree of synchrony in their CDT1a loading pattern.

**Figure 2.**
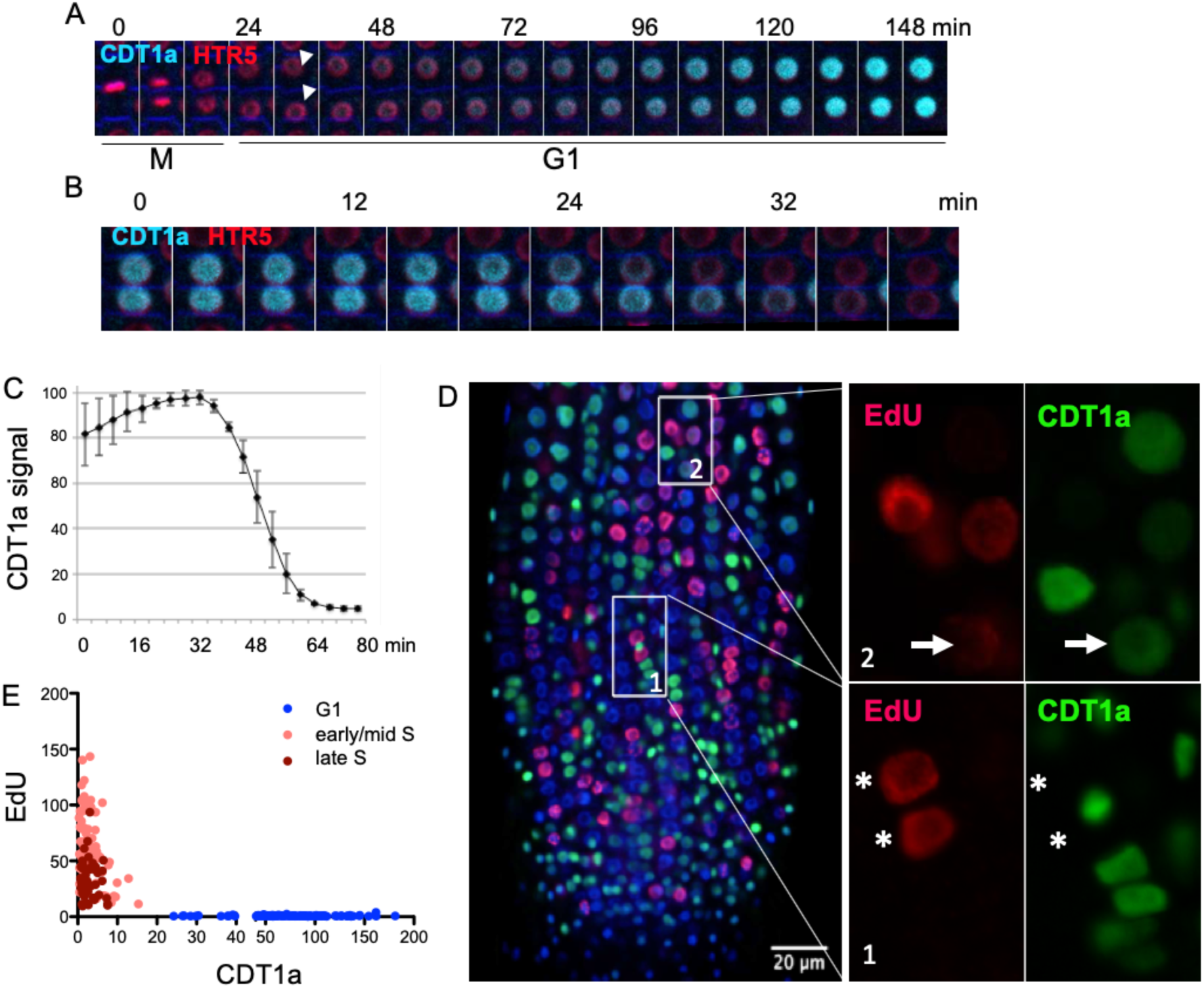
Dynamics of CDT1a during the cell cycle. (A) Live imaging visualizing CDT1a loading during G1. The arrowheads mark two newly-formed daughter cells entering the G1 phase. CDT1a-CFP (cyan) level is shown at different time points, as indicated. Nuclei were detected by the constitutive expression of histone H3.3 HTR5-mRFP (red). (B) Live imaging showing the degradation of CDT1a-CFP (cyan). All nuclei are visualized as in (A) with HTR5-mRFP (red). Two cells are shown to illustrate the fast CDT1a degradation process. (C) Quantification of the CDT1a degradation kinetics by measuring the CFP signal intensity. Quantification was performed from cells (N = 4) exhibiting a CDT1a-CFP signal from the mid/late G1 until the complete loss of the cyan signal in the analyzed cell. Values were normalized to 100% at maximum fluorescence value. (D) Regulation of CDT1a-CFP during S-phase. Roots were labeled for 15 minutes with EdU (red) whereas CDT1a (green) was detected by immunofluorescence and all nuclei by DAPI staining (blue). Insets illustrate cases where the EdU+ nuclei are devoid of CDT1a (inset 1) and where a faint EdU and CFP signals colocalize (arrows in inset 2). (E) Quantification of CDT1a and EdU signals in nuclei with the indicated late or early/mid EdU labeling pattern. Note that these two signals are largely excluded from the same nucleus. Quantification was performed from nuclei (N≥200) in the RAM zone of the above described material (D).

Live imaging also revealed that nuclei containing the highest levels of CDT1a-CFP eventually lose the CDT1a-CFP signal (Figure 2B–movie supplement 1). The degradation kinetics was very fast (∼20-30 min to reach undetectable levels of CDT1a), as demonstrated by quantifying the loss of CDT1a-CFP signal (Figure 2C).

To find out when during the cell cycle degradation of CDT1a occurs we first asked whether proliferating cells containing CDT1a-CFP are progressing through S-phase by labeling with a 15 min pulse of 5-ethynyl-2’-deoxyuridine (EdU). We found that CDT1a-CFP and EdU signals did not colocalize in almost any of the nuclei analyzed (Figure 2D, inset 1). In the few cases found with detectable amounts of both CDT1a-CFP and EdU (<2%), the signals were very low (Figure 2D, inset 2). Independent quantification of the EdU and CDT1a signals further confirmed that most nuclei contained either signal but not both (Figure 2E). Furthermore, colocalization of both signals corresponded with an early S-phase EdU labeling pattern (Dvorackova et al., 2018), supporting the conclusion that CDT1a-CFP is degraded upon entering S-phase. Therefore, we conclude that CDT1a is deposited early after mitosis and is degraded rapidly as soon as cells pass the G1/S transition.

### CDT1a is loaded immediately after S-phase in endoreplicating cells

Cells undergoing the endocycle in the transition zone of the root must replicate their genome during S-phase. However, the dynamics of pre-RC proteins, our focus here, as well as other mechanistic aspects of the endocycle S-phase are not well established. It is known that Cdt1 oscillates both in mitotic and endocycling Drosophila cells and exhibits low levels during S-phase (Thomer et al., 2004; Whittaker et al., 2000). We found that in the transition zone of the root where cells are engaged in the endocycle, more cells contained CDT1a than in the proliferative area but the pattern was also patchy with cells showing various levels of CDT1a-CFP, including a fraction of nuclei devoid of the protein (Figure 3A). A similar pattern was observed in the *cdt1a-1* plants complemented with the CDT1a-CFP fusion protein (Figure 3B). To determine the CDT1a status in relation to the endocycle S-phase we labeled with EdU and found that the vast majority of cells in the transition zone did not have detectable levels of CDT1a-CFP in EdU-labeled cells (Figure 3C, inset 1), and only a small amount showed low levels of CDT1a-CFP and EdU (Figure 3C, inset 2). This pattern was similar to that of the meristematic cells and strongly indicative of a fast degradation of CDT1a-CFP upon S-phase initiation during the endocycle. Therefore, we can conclude that the CDT1a dynamics during S-phase of both proliferating and endocycling cells is similar.

**Figure 3.**
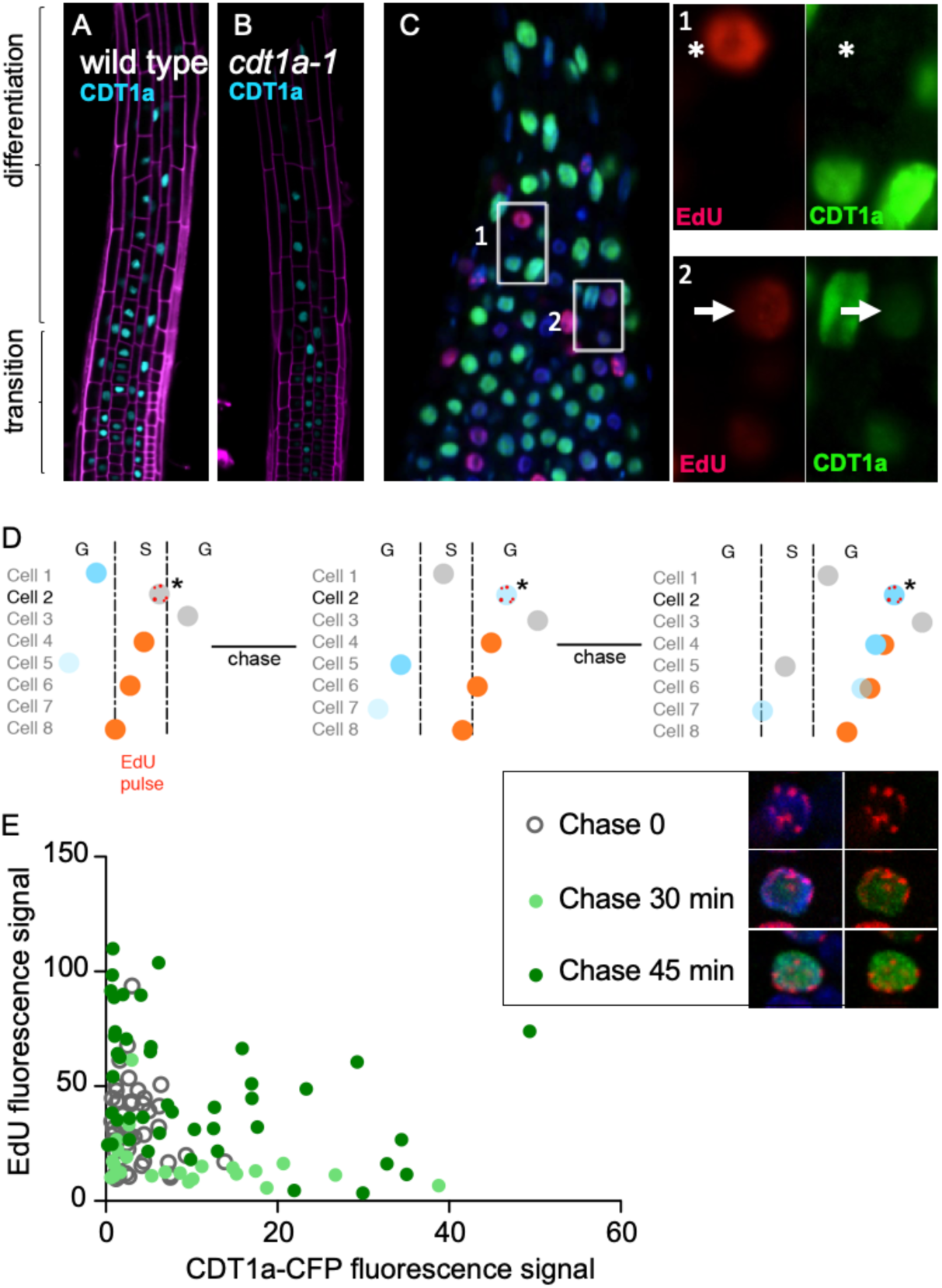
Dynamics of CDT1a in endoreplicating cells. (A,B) Expression of CDT1a-CFP in endoreplicating cells of the transition zone of the root apex in a wild type (A) and in the *cdt1a-1* (B) mutant backgrounds. (C) CDT1a dynamics during the endocycle G phase. Most cells possess the pattern shown in inset 1, where EdU-labeled nuclei are devoid of CDT1a. Example of cells exhibiting a low signal of CDT1a colocalizing with a low EdU signal (inset 2). All nuclei were counterstained with DAPI (blue). (D) Schematic of the strategy to visualize CDT1a loading during the G phase of the endocycle. Cells were pulse-labeled with EdU (15 min). Quantification of the time of appearance of CDT1a signal in EdU-labeled nuclei with a late-replicating pattern (labeled chromocenters; cell #2*) at different chasing times gives a direct measurement of CDT1a incorporation after the endocycle S-phase. (E) Quantification of EdU and CDT1a-CFP signals at different chasing times of nuclei with a late-replicating EdU-labeling pattern (N=48, N=22, N=42 nuclei for chasing times 0, 30 and 45 min, respectively).

After termination of the endocycle S-phase, cells skip mitosis and enter a G phase before a new endoreplication round initiates. A key question is when after S-phase of endocycling cells start to load again CDT1. To answer this, we labeled root cells with EdU (15 min) and chased them for various times. The time taken by EdU-labeled cells with a late-replicating pattern (labeled chromocenters, Fig. 3D) to show a detectable CDT1a-CFP signal is a direct measurement of the time required for CDT1a to load during the endocycle G phase after S-phase termination. We found that CDT1a is loaded just after a short (∼20-30 min) time in late-replicating nuclei (Figure 3E). Our results demonstrate that the G phase of the endocycle behaves largely as a G1 phase, based on the CDT1a dynamics.

### Role of CDT1a domains in its dynamics and genome stability

CDT1 is not highly conserved in sequence except for a ∼150 amino acid-long C-terminal domain, involved in MCM interaction (Castellano et al., 2004; Ferenbach et al., 2005). We confirmed the relevance of CDT1 domains by testing whether two truncated versions of CDT1a were able to complement the lethal *cdt1a-1* mutation (Figure 5A). We found that expression of the N-terminal moiety (N-CDT1a; amino acids 1-362) was unable to rescue viability of *cdt1a-1* mutant (n=52 plants genotyped), in spite of having the expected expression domain (Figure 4B), but consistent with previous studies with animal Cdt1 (Ferenbach et al., 2005). On the contrary, expression of the C-terminal moiety (C-CDT1a; amino acids 363-571), a truncation proposed recently as a cell cycle marker in pants (Yin et al., 2014), did allow the recovery of viable plants, allowing further studies (Figure 4G). The expression domain of the C-CDT1a in the RAM, both in a wild type and a *cdt1a-1* mutant background, was different from the full-length protein (see also Figure 1 and 3), as it was present in many proliferating cells in the RAM as well as in the transition zone of the root but not in the elongation zone (Figure 4C,D). Importantly, this truncated version did not show the same dynamics during the cell cycle as the full-length protein, as revealed by an EdU pulse, showed S-phase cells containing high levels of C-CDT1a (Figure 4E), consistent with previous reports (Yin et al., 2014). Since the full-length protein is rapidly degraded upon S-phase initiation, the maintenance of the C-terminal moiety during S-phase allowed us to evaluate whether it had deleterious effects. Indeed, flow cytometry analysis revealed that excess of C-CDT1a in a *cdt1a-1* background during S-phase led to an abnormal nuclear ploidy profile with clear evidence of partially replicated genomes whereas expression of C-CDT1a in wt and expression of full length CDT1a in *cdt1a-1* showed comparable ploidy profiles (Figure 4F). Since the C-terminal moiety bears the MCM interaction domain, the effect observed could be a consequence of altered availability of MCM, strongly suggesting that elimination of CDT1a early upon S-phase entry is a mechanism to prevent genome instability. Moreover, plants expressing the C-CDT1a in the *cdt1a-1* mutant background could grow well but they reached smaller size (Figure 4G), produced smaller siliques (Figure 4H) and contained aborted seeds (Figure 4I).

**Figure 4.**
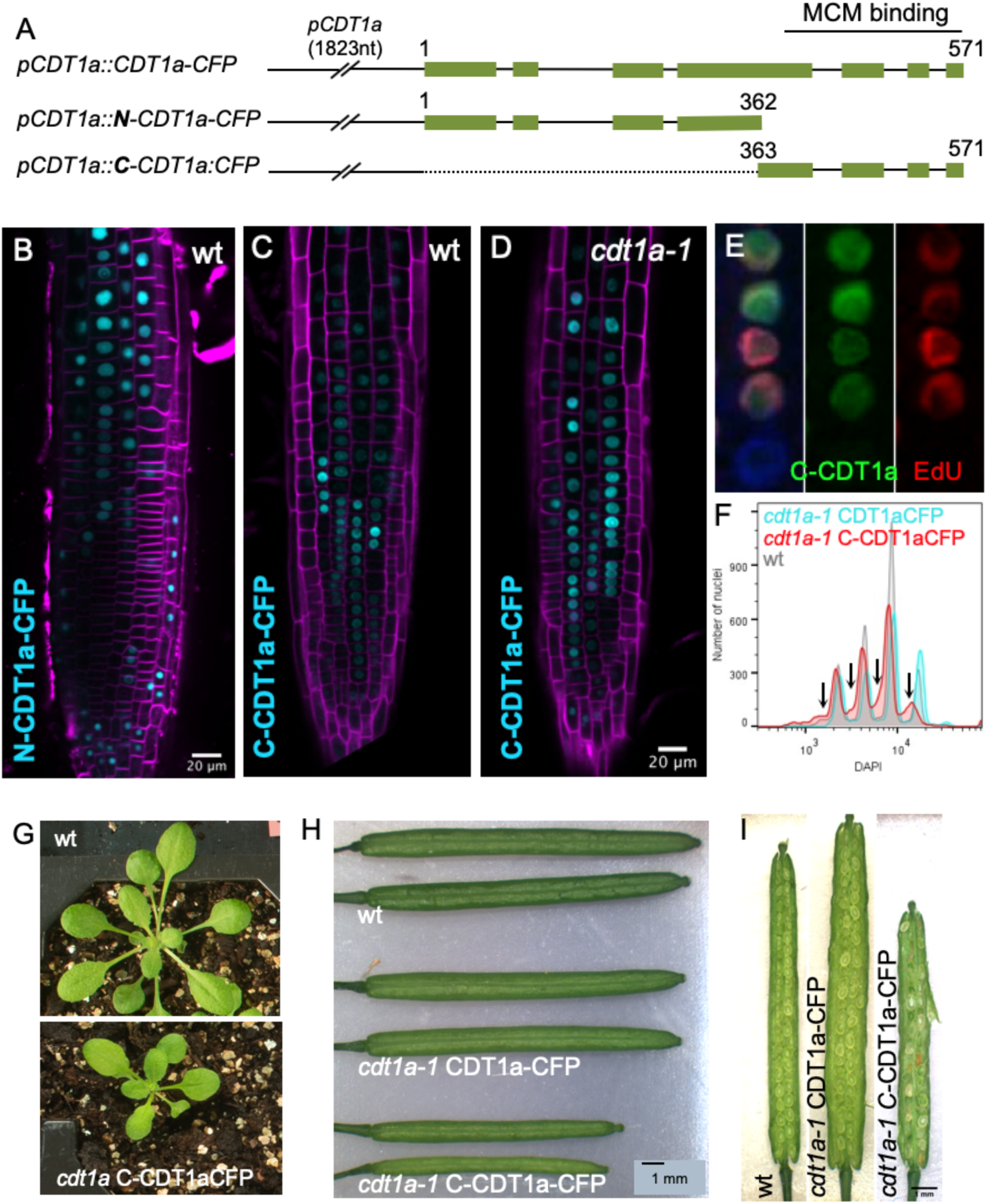
Role of CDT1a domains in degradation and genome instability. (A) Schematic of the CDT1a full-length and deletion constructs expressed under the endogenous *pCDT1a* promoter. The MCM binding domain is located at the C-terminus of CDT1a. Note that the construct expressing the N-terminal moiety of CDT1a was not able to complement the *cdt1a-1* lethal phenotype. (B) Expression of the N-terminal moiety (N-CDT1a-CFP) in wild type plants. (C) Expression of the C-terminal moiety (C-CDT1a-CFP) in wild type plants. (D) Expression of the C-terminal moiety (C-CDT1a-CFP) in *cdt1a-1* mutant plants. (E) The C-CDT1a truncated protein is not degraded at the onset of S-phase, as it occurs with the full-length CDT1a protein. (F) Representative nuclear ploidy profile of wild type plants (grey) and *cdt1a-1* mutants plants expressing the full length CDT1a (cyan) or the C-terminal moiety (C-CDT1a; red). Note the abnormal profile of these nuclei with increased amounts of partially replicated genome (arrows). Leaves #3/4 of 22-day-old plants were used. The experiment was repeated at least 3 times with similar results. (G) Complementation of the *cdt1a-1* mutation with the CDT1a C-terminal half (C-CDT1a). (H) Siliques of control (wt) and transgenic plants expressing the C-CDT1a in the wild type or *cdt1a-1* mutant background. (I) Detail of siliques shown in panel H to highlight the presence of aborted seeds.

**Figure 5.**
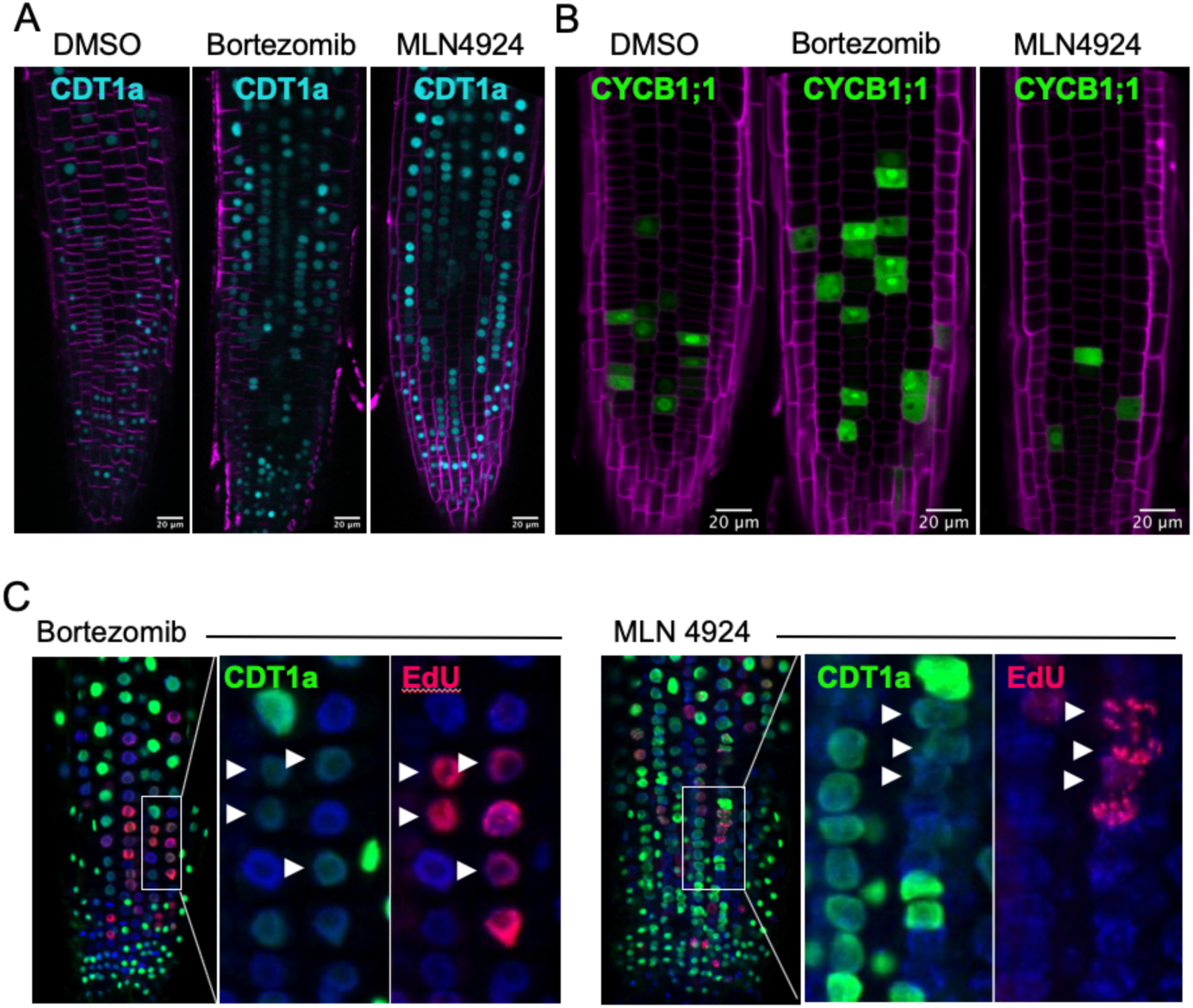
CDT1a is targeted for proteasome degradation. (A) The expression of CDT1a-CFP (cyan) was assessed by confocal microscopy of root apices after treatment with DMSO (control) and proteasome inhibitors bortezomib (50 µM) or MLN4924 (25 µM) for 2.5 hours. (B) Effect of bortezomib and MLN4924 as in panel (A) on the CYCB1;1-GFP protein. (C) Detection of S-phase progression in the presence of proteasome inhibitors, as indicated. Roots were labeled with EdU (30min; red) at the end of the treatment with the indicated proteasome inhibitors. CDT1a-CFP was detected by immunofluorescence (green). All nuclei were counterstained with DAPI. Insets show details of the indicated panels. Arrowheads point to nuclei in S-phase also containing CDT1a-CFP.

### CDT1a is targeted for proteasome degradation

Degradation of Cdt1 in animal cells occurs through the proteasome (Havens and Walter, 2011; Nishitani et al., 2006). To assess whether this degradation pathway appeared early in evolution we treated Arabidopsis seedlings expressing CDT1a-CFP with bortezomib, an inhibitor of the catalytic subunit of the 26S proteasome (Bonvini et al., 2007). Treatment led to the accumulation of CDT1a-CFP in meristematic cells (Figure 5A). As a control we also treated seedlings expressing CYCB1;1-GFP (Ubeda-Tomas et al., 2009) which is degraded by the APC in anaphase. As expected, bortezomib-treated cells showed a significant accumulation of cells in anaphase with high amounts of CYCB1;1 (Figure 5B). To get further information, we treated plants with bortezomib during 4 h and pulsed-labeled them with EdU during the last 30 minutes of the treatment. We found that CDT1a-CFP colocalized with EdU after the bortezomib treatment, indicating that it was no longer degraded at the beginning of S-phase and that cells can proceed into S-phase with high levels of CDT1a-CFP.

Similar accumulation of CDT1a was observed with another proteasome inhibitor, MLN4924, which prevents the neddylation step required for SCF activity (Figure 5A), revealing that CDT1a proteasome targeting requires ubiquitylation by an SCF-type E3 ligase. Since the neddylation step is not required for the APC/C activity, another type of E3 ligase, MLN4924 does not affect stability of CYCB1;1 (Figure 5B). Again, cells can be identified containing both CDT1a-CFP and EdU signals (Figure 5B).

### FBL17 targets CDT1a for proteasome degradation

To identify the mechanism of targeting CDT1a for proteasome degradation we crossed CDT1a-CFP-expressing plants with several known mutants defective in cell cycle and growth-related SCF function. We found that both the expression pattern and the amount of CDT1a-CFP per nucleus were very similar in the *skp2a, cul4a* and *skp2a,cul4a* mutant backgrounds compared to controls (Figure 6A). This result was confirmed by western blot analysis that showed no increased levels of CDT1a-CFP in all mutants analyzed (Figure 6B). Therefore, we conclude that although CDT1a is likely targeted for degradation by a SCF complex soon after the onset of S-phase, neither SKP2A nor a CRL-type of ubiquitin E3 ligase is involved in CDT1 degradation.

**Figure 6.**
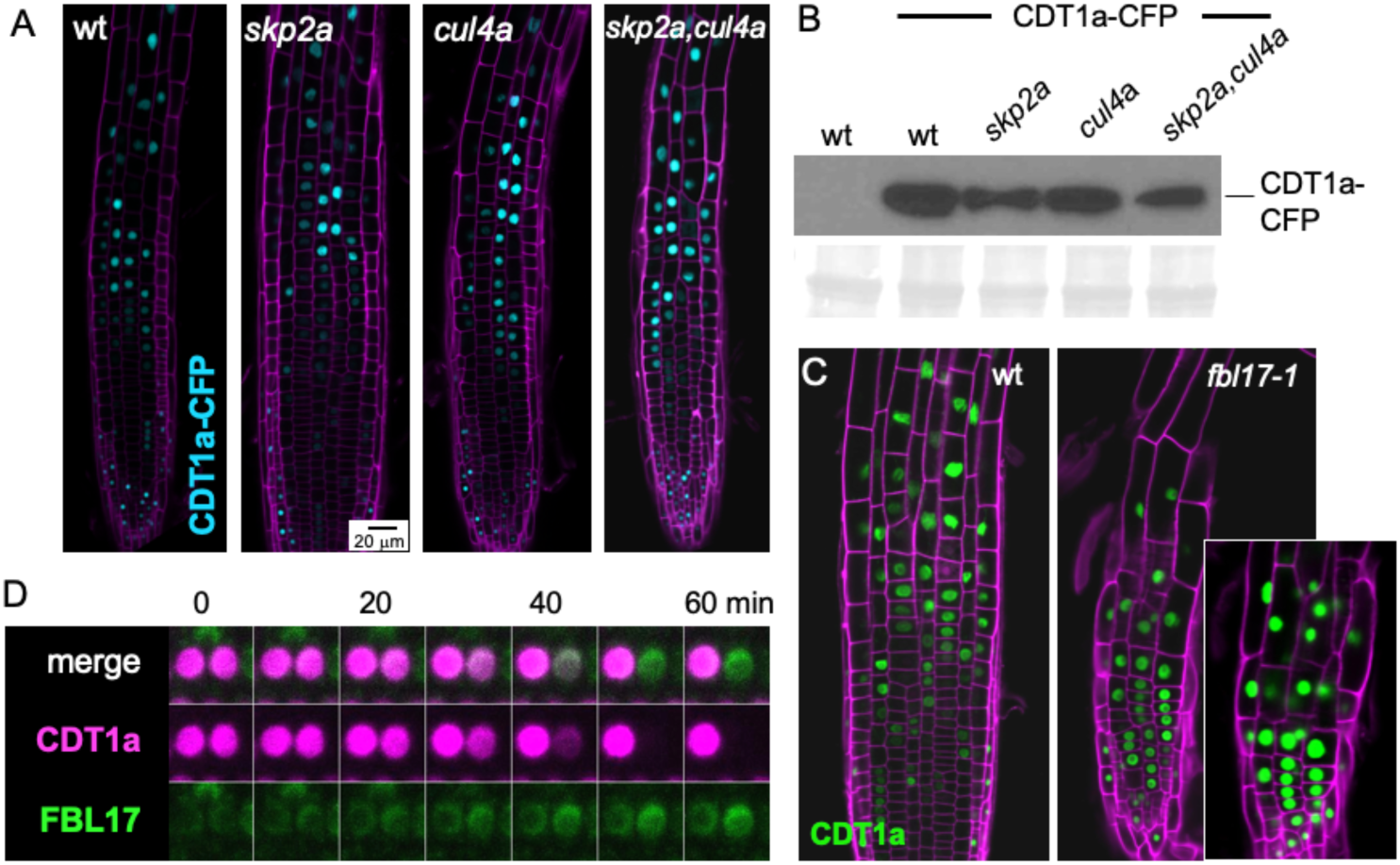
Identification of FBL17 as a F-box protein involved in CDT1a degradation. (A) Expression of CDT1a-CFP (cyan) assessed by confocal microscopy in root tips of wild type and indicated mutants of the proteasome pathway. Cell walls were detected by staining with propidium iodide (magenta). (B) Western blot revealing the amount of CDT1a-CFP protein expressed in the indicated lines. (C) Expression of CDT1a-CFP assessed by confocal microscopy in wild type and *fbl17-1* mutant root tips. The saturated image (inset) was acquired with the same confocal setting as in panel C (left). Cell walls were detected as in panel A. (D) Live imaging showing the dynamics of CDT1a-mRFP (magenta) and FBL17-GFP (green) during the G1 and early S-phase. Note that the CDT1a signal disappears rapidly as the FBL17 protein reaches a certain threshold.

Among the very large Arabidopsis family of F-box proteins, FBL17 is a key factor involved in cell cycle transitions during vegetative growth as well as during cell divisions of the germline (Gusti et al., 2009; Kim et al., 2008; Noir et al., 2015; Zhao et al., 2012). Thus, we hypothesize that FBL17 is a good candidate to target CDT1a for proteasome degradation. To test this, we crossed plants expressing CDT1a with heterozygous *fbl17-1*^*+/-*^ plants. In spite that the *fbl17-1* mutation is lethal, ∼1% of the embryo progeny can be rescued and grow to produce an abnormal seedling (Noir et al., 2015). Although with a very altered root meristem it was clear that CDT1a-GFP accumulates and most cells maintain high levels of the protein in the *fbl17-1* mutant background, compared to the control plants (Figure 6C). To further assess the involvement of FBL17 in CDT1a degradation we determined the dynamics of both proteins in plants expressing CDT1a-mRFP and FBL17-GFP. Live imaging demonstrated that high levels of CDT1a are maintained in nuclei while FBL17 protein starts to accumulate and then rapidly diminished once the maximum levels of FBL17 are reached (Figure 6D and movie supplement 2).

### FBL17-mediated CDT1a degradation is robust independently of protein levels

It has been reported that the *fbl17-1* mutant shows altered expression of several cell cycle regulatory proteins (Noir et al., 2015). We confirmed that both *CDT1a* and *E2Fa* are upregulated in *fbl17-1* seedlings (Figure 7A). Since E2Fa expression is up-regulated in *fbl17-1* mutants and *CDT1a* is transcriptionally activated by E2Fa (Castellano et al., 2004; Ramirez-Parra et al., 2003; Vlieghe et al., 2005) one possibility is that the excess CDT1a levels detected in the *fbl17-1* mutant is solely a consequence of transcriptional upregulation by the increased levels of E2Fa. To test this possibility, we generated plants expressing CDT1a under the control of a constitutive promoter, which is not regulated by E2Fa. Western blot confirmed the high amount of CDT1a expressed (Figure 7B). The protein is loaded in G1 and degraded at the beginning of S-phase (Figure 7–figure supplement 3A), indicating that CDT1a is regulated at the posttranscriptional level in the meristem also under constitutive expression. CDT1a-CFP accumulates at high levels in the differentiated zone and endoreplication is enhanced (Figure 7–figure supplement 3B,C). We also selected by crossing plants expressing CDT1a-CFP constitutively in the *fbl17-1* mutant background and found that CDT1a accumulates in most proliferating cells of the small meristem of the *fbl17-1* mutant (Figure 7C), but to a lesser amount than to when CDT1a expression is controlled by its own promoter. Together these results indicate that the accumulation of CDT1a in the *fbl17-1* background is due to a combination of both inhibition of selective proteolysis and transcriptional upregulation. Importantly, we found that mid S-phase cells labeled with EdU contained a significant amount of CDT1a (Figure 7D,E) indicating that indeed FBL17 is necessary to drive CDT1a degradation at the initiation of S-phase. Thus, we conclude that CDT1a regulation, including accumulation during G1 and rapid degradation early upon S-phase entry, is very robust at the protein level, independently of the mRNA or protein expression levels.

**Figure 7.**
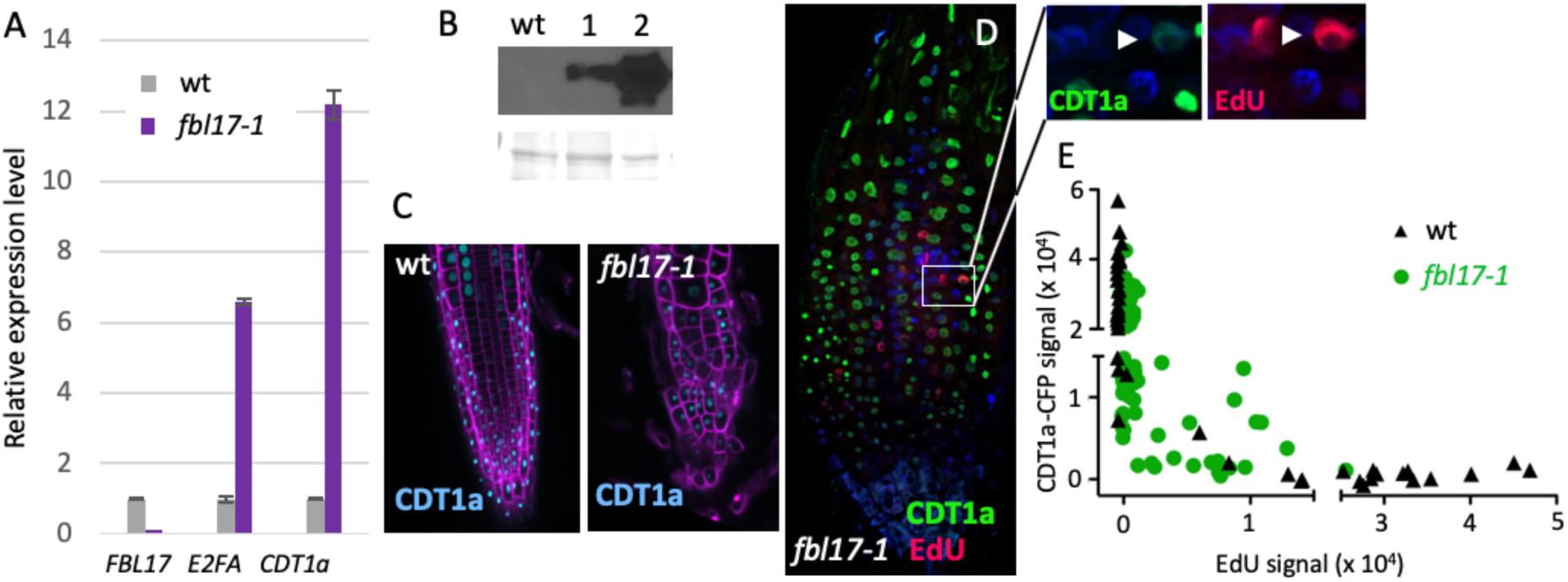
Regulation of CDT1a levels expressed under a constitutive promoter. (A) Relative expression levels of *FBL17, E2FA* and *CDT1a* transcripts were determined by qRT-PCR in wild-type and *fbl17-1* seedlings. The bar graph depicts the mean values of the indicated transcripts of one independent replicate (± standard errors of the technical triplicate). The experiment was repeated at least 3 times giving similar differential expression levels. (B) Western blot revealing the accumulation of CDT1a-CFP in plants expressing CDT1a from its own promoter (lane 1) and from a constitutive promoter (lane 2). Loading control is shown at the bottom. (C) CDT1a dynamics in wild type and *fbl17-1* mutant, expressed with a constitutive promoter. (D) Detection of S-phase progression and CDT1a levels under constitutive expression in the *fbl17-1* mutant background. Roots were labeled with EdU (30 min; red). CDT1a-CFP was detected by immunofluorescence (green). Insets show details of the indicated panel. All nuclei were stained with DAPI. (E) Quantification of EdU and CFP signal intensities in nuclei of plants expressing constitutively CDT1a-CFP in wt (black triangles) and *fbl17-1* (green circles) mutant backgrounds (N=76, N=52 nuclei for wild type and *fbl17-1* mutant, respectively).

## Discussion

A strict control of chromosomal replication is crucial to maintain genome stability. A primary step that is tightly controlled is the initiation of DNA replication that must occur once and only once per cell cycle. Thus, cells have evolved mechanisms to prevent that DNA replication origins (ORIs) are activated more than once in the same S-phase, so-called re-replication, by sequential degradation of key factors (Arias and Walter, 2007; Coleman et al., 2015). To prevent this harmful scenario for genome stability it is necessary to regulate the availability and function of key factors acting at the early stages of ORI activation after assembly of pre-replication complexes (pre-RC). However, sufficient amounts of chromatin-bound pre-RCs are necessary for correct full genome replication because limiting amount of pre-RC or ORI activation in excess cause partial replication phenotypes (Blow et al., 2011; Moreno et al., 2016). Several pre-RC components are subject to control their nuclear abundance (Asano and Wharton, 1999; Castellano et al., 2001; DePamphilis, 2003; Kim and Kipreos, 2008; Kuo et al., 2012; Ohtani et al., 1998), but Cdt1 is probably the most tightly regulated pre-RC component (Arias and Walter, 2007; Castellano et al., 2004; Dorn et al., 2009; Thomer et al., 2004). Cdt1 abundance needs to be high to ensure sufficient pre-RC assembly and ORI activation but also its nuclear level has to be reduced at some point after ORI activation to prevent re-replication. Thus, overlapping, although not redundant mechanisms operate at levels of transcriptional regulation, protein stability and activity, and involve species-specific mechanistic differences (Arias and Walter, 2007).

Control of Cdt1 abundance is first exerted at the transcriptional level both in unicellular and multicellular eukaryotes. In *S. pombe* the CDC10 transcription factor, which also regulates other cell cycle genes (Whitehall et al., 1999), drives the production of high Cdt1 mRNA levels (Hofmann and Beach, 1994). In animal and plant cells, transcription of Cdt1 gene depends on the E2F family of transcription factors and the retinoblastoma-related proteins (Castellano et al., 2004; Desvoyes et al., 2006; Ramirez-Parra et al., 2003; Yoshida and Inoue, 2004). Consistent with this, we found progressive accumulation of a detectable signal of Arabidopsis CDT1a protein shortly after initiation of G1, which reached a maximum late in G1. Therefore, this mechanism appears to be conserved, although with different players in unicellular (yeast) and multicellular (animals, plants) organisms.

A second pathway functioning in animal cells is the inhibition of Cdt1 activity by geminin, a protein that is regulated by the E2F pathway (Markey et al., 2004; Yoshida and Inoue, 2004), increases in S-phase and is maintained at high level until anaphase where it is degraded by the APC (McGarry and Kirschner, 1998). Geminin regulates DNA replication licensing (DePamphilis et al., 2006) by interacting with Cdt1 and inhibiting MCM loading (Maiorano et al., 2004; Wohlschlegel et al., 2000). Interfering with geminin abundance in animal cells by altering its expression or proteolysis leads to genome instability (Neelsen et al., 2013; Zhu et al., 2004). Geminin appears to be an evolutionary acquisition of the animal lineage since it is not present in plant genomes, in spite that Arabidopsis encodes GEM, a CDT1-interacting protein, with remarkable functional analogies with animal geminin, but structurally unrelated (Caro et al., 2007; Caro and Gutierrez, 2007; Kroll, 2007; Luo et al., 2004; Seo and Kroll, 2006). However, any potential role of GEM in CDT1a-related DNA replication functions has not been addressed yet.

Proteasome-mediated degradation of Cdt1 seems to be a third mechanism in animal as well as in plant cells as demonstrated in this work. This is different from yeast where Cdt1, in complex with MCM, is exported out of the nucleus during S-phase (Nguyen et al., 2000; Tanaka and Diffley, 2002). Two different degradation pathways have been demonstrated in animal cells, mediated by the Skp2 and Cdt2 CRL receptors (Havens and Walter, 2011; Nishitani et al., 2006). Similar to animal Cdt1 that requires a SCF^Skp2^ complex, Arabidopsis CDT1a is targeted for degradation by a SCF^FBL17^ complex, a process that likely requires CDK activity (Castellano et al., 2004). The other pathway that controls human Cdt1 level is mediated by the Cdt2 protein, which is part of a CRL4 complex and instead of phosphorylation, requires interaction of Cdt1 with PCNA (Hayashi et al., 2018; Mazian et al., 2019; Nishitani et al., 2006). This Cdt1 targeting mechanism is mediated by a PCNA-interacting protein (PIP) motif located at the N-terminus of human Cdt1 (Havens and Walter, 2011), which is not conserved in all animal species. The lack of a PIP box in Arabidopsis CDT1a makes unlikely that this pathway operates in plants to regulate CDT1a abundance.

The plant F-box family contains >700 members (Gagne et al., 2002; Lechner et al., 2006). Among them, FBL17 has been reported as a crucial factor controlling cell proliferation in vegetative and germline tissues, endoreplication in various organs and, more specifically, the G1/S transition (Gusti et al., 2009; Kim et al., 2008; Noir et al., 2015; Zhao et al., 2012). Therefore, FBL17 appeared as an excellent candidate to be part of an SCF complex targeting CDT1a for degradation. Indeed, the *fbl17-1* mutant shows a major accumulation of CDT1a, which may be at the basis of its defects in genome stability and cell proliferation (Gusti et al., 2009). It seems that a distinct F-box protein has evolved in plants to regulate the abundance of CDT1a as a mechanism for genome maintenance. Another difference with the regulatory mechanism acting on Cdt1 in animals (Rizzardi et al., 2015) is the absence of Arabidopsis CDT1a during the G2 phase. Although we cannot discard the existence of other degradation pathways, it seems that Arabidopsis CDT1a abundance is mainly regulated at the transcriptional level and by selective proteolysis. Given the importance of regulating DNA replication by overlapping mechanisms to prevent genome instability it is conceivable that additional safeguard mechanisms could have evolved in plants acting on other pre-RC components. The rationale and the tools available now should facilitate addressing these questions in the future to gain insight into the evolutionary history of the mechanisms preventing genome instability in multicellular eukaryotes.

## Materials and methods

### Plant growth

After stratification of *Arabidopsis thaliana* (Col-0 ecotype) seeds for 48h, plants were grown in 0.5x MS medium (pH 5.7) supplemented with MES, vitamins, 0.5 or 1% sucrose and 1% plant agar (Duchefa) at 21 °C and 60% moisture, under long day conditions (16 h light, 8 h dark). MS medium was supplemented with antibiotics for plant selection and the same MSS medium without agar was used for metabolic labeling and drug treatment.

### Generation of CDT1a reporter lines

For the pCDT1a:CDT1a-CFP, pCDT1a:CDT1a-GFP and pCDT1a:CDT1a-mRFP constructs, the genomic fragment encompassing 1823bp of the promoter and the 2505 bp of the gene was cloned into the pDONOR221 using the gateway system and then inserted in frame with the FP into the destination vector pGWB443 for fusion with eCFP, pGWB450 for GFP and pGWB453 for mRFP (Nakagawa T. et al., 2009). The C-CDT1a-CFP was generated by PCR mediated deletion of the 1578 bp N-terminal region using as a template the pCDT1a:CDT1a entry clone (Hansson et al., 2008). For the N-CDT1aCFP construct, the 1823 bp CDT1a promoter and the 1577 bp 5’ moiety of gCDT1a gene were amplified by PCR using the KOD polymerase (Millipore) and domesticated into the pUPD2. The different modules (promoter, gene, eCFP and HSP18 terminator) were then assembled into a binary vector using the Goldenbraid system (Sarrion-Perdigones et al., 2014). The Goldenbraid system was also used to generate the construct for the CDT1a ubiquitous expression line, using the pU6 promoter (At3g13855, 388bp), the genomic fragment of CDT1a coding sequence, eCFP and HSP18 terminator. Primers used in this study are listed in Supplementary Table 1 (relate to Figures all figures). All constructs were checked by sequencing (Macrogen). Transgenic plants were generated by the floral dip method (Clough and Bent, 1998), using the *Agrobacterium tumefaciens* C58C1 strain and transformants were selected with kanamycin.

The *cdt1a-1* GABI_KAT_025G08 knockout line was described earlier (Domenichini et al., 2012) and obtained from the NASC. Mutants of the proteasome pathway *fbl17-1* (GABI_KAT_170E02), *cul4-1, skp2a* (GABI_KAT_293D12), *skp2b* (SALK_028396) have also been described previously (Bernhardt et al., 2006).

For genotyping the T-DNA insertion mutants, total genomic DNA was extracted. One frozen leaf was grinded, resuspended in extraction buffer (d-Sorbitol 0.14M, Tris-HCl pH8 220 mM, EDTA 22 mM, NaCl 0.8 M, CTAB 0.8%, n-laurylsarcosine 1%) incubated at 65°C for 10 min and extracted with chloroform. DNA was precipitated with ethanol and resuspended in water. PCR amplification was carried out using primers listed in Supplementary Table 1 (relate to all figures) and Taq polymerase (NZYTech).

### RNA extraction and RT-qPCR

Purification of total RNAs from 12-day-old seedlings grown under *in vitro* conditions was performed using the Tri-Reagent (Molecular Research Center, Inc.) according to the manufacturer’s instructions. cDNAs were prepared using the High Capacity cDNA Reverse Transcription kit (Applied Biosystems™). Real-time amplification was carried out using gene-specific primers and SYBR Green Master mix (Roche) on a Lightcycler LC480 apparatus (Roche) according to the manufacturer’s instructions. The mean value of three replicates was normalized using the *EXP (AT4G26410)* and *TIP41 (AT4G34270)* genes as internal controls. Primers used in quantitative RT-PCR are listed in Supplemental Table 1.

### EdU labeling

Four-day-old seedlings were incubated in liquid MSS supplemented with 20 µM of EdU for 15 or 30 min at room temperature and when indicated, washed once with MSS and chased for the specified periods of time with 50 µM thymidine. Plants were then fixed with paraformaldehyde 4% and immunofluorescence was performed as described previously (Otero et al., 2016). Roots were imaged by confocal microscopy using a Zeiss LSM 710 or LSM800.

### Confocal microscopy

Five- or 6-day-old roots were stained with propidium iodide (50 µg/mL) and observed using a Zeiss LSM710 confocal microscope. For live imaging, 4-day-old plants were transferred to a P35 glass bottom dish, covered with a piece of 1% agar MSS and acclimated for 4-10h in the growth chamber. FM4-64 (10 µM) was used to stain plasma membrane and images were acquired with a NIKON A1R+ confocal microscope. Quantification of fluorescence signal and movie editing were performed using FIJI.

### Flow cytometry

Leaves #1-2 and #3-4 of 3 week-old plants were chopped with a razor blade in 500 µl of nuclear isolation buffer (Desvoyes et al., 2006). DAPI was added at 2 µg/ml and nuclei were analyzed using a FACSCanto II flow cytometer (Becton Dickinson).

### Protein extraction and Western blot analysis

Nuclear extract was prepared with 0.2 g of seedlings (5 days after sowing) essentially as described previously (Desvoyes et al., 2018). The nuclear lysate was subjected to sonication in a Bioruptor (Diagenode), 10 cycles 30’ On, 30’ Off. After centrifugation the cleared extracts (15 µg of proteins) were fractionated in a 10% polyacrylamide gel and protein immunodetection was realized with an antibody anti GFP (PABG1, Chromotek, 1/3000).

## Acknowledgments

We thank the Confocal microscopy service of CBMSO for support, E. Martinez-Salas for comments, and members of the laboratory for continuous feedback. Research in our laboratories was supported by grants BFU2015-68396-R, BIO2017-92329-EXP, RTI2018-094793-B-I00 and grant ERC-2018-AdG_833617 to C.G., grant LABEX ANR-10-LABX-0036-NETRNA to P.G., and by institutional grants from Banco de Santander and Fundación Ramon Areces to the CBMSO.

## Author contributions

B.D. and C.G. designed the research with the contribution of P.G. B.D. and S.N. performed experiments with the initial contribution of K.M. and M.I.L. All authors analyzed and interpreted the data. C.G. and B.D. wrote the manuscript with the input of P.G. and S.N. All authors revised and agreed the content of the manuscript.

## Figure supplements

**Figure S1.**
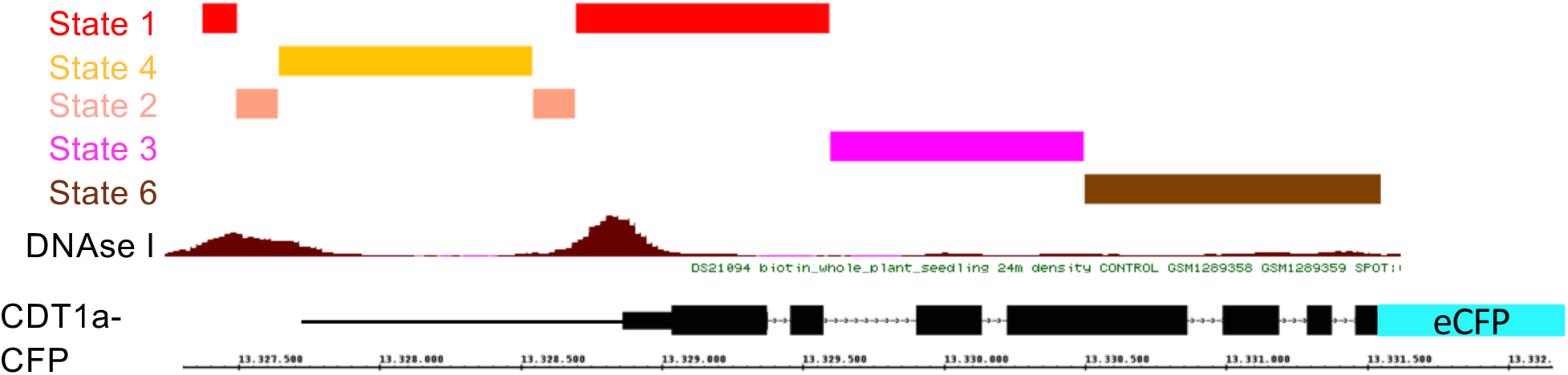
Gene organization and chromatin landscape of the CDT1a genomic region. The different chromatin states, as described (Sequeira-Mendes et al., 2014) covering the gene as well as the upstream region are indicated together with the DNase I hypersensitive sites. The color code is red (state 1, active gene), salmon (state 2, proximal promoter), pink (state 3, transcribed gene body), yellow (state 4, intergenic region) and brown (state 6, 3’ of transcribed gene).

**Figure S2.**
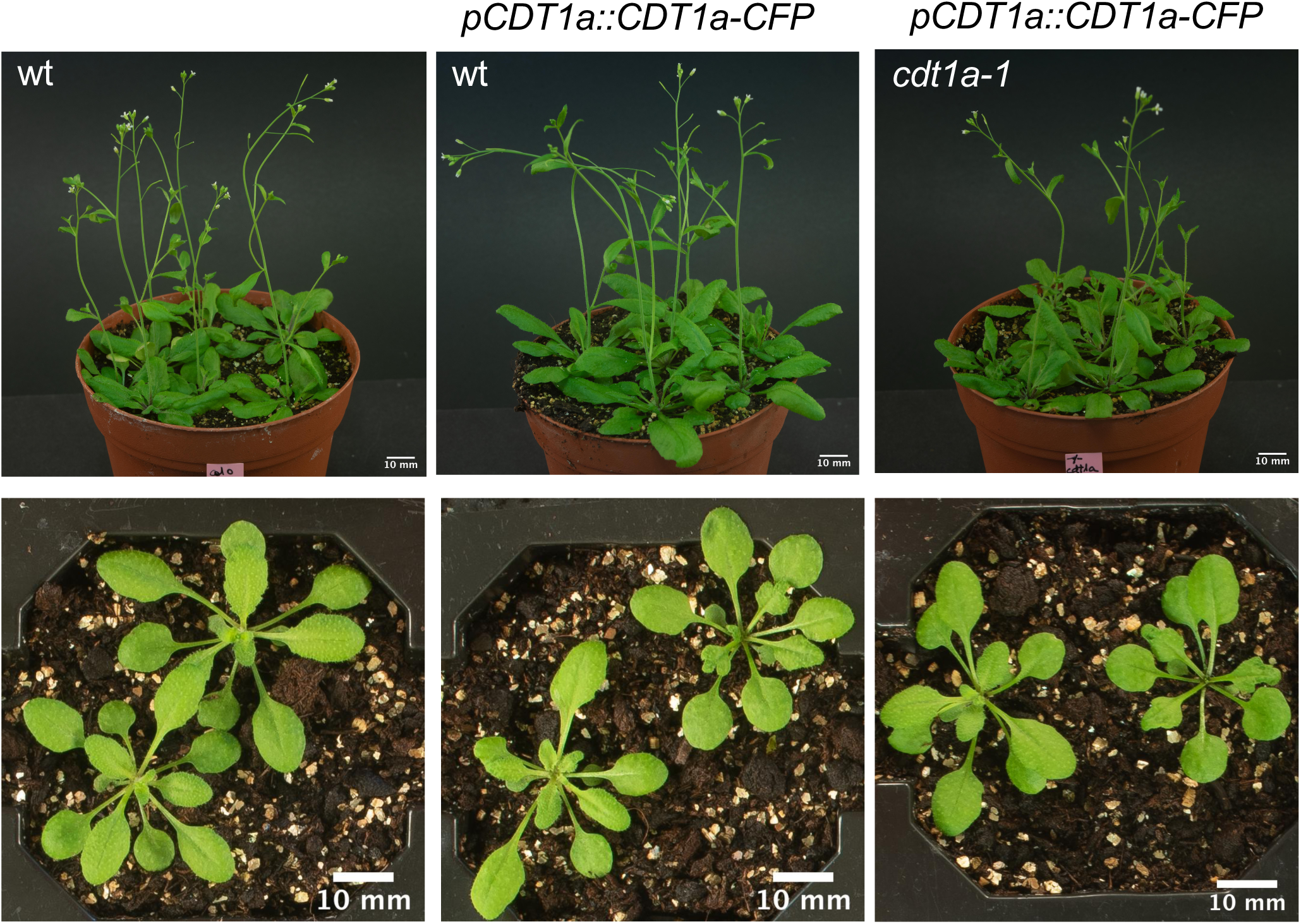
Phenotypic analysis of plants expressing the CDT1a-CFP fusion protein. The construct expressing CDT1a-CFP under its own promoter was expressed in wild type (wt) and *cdt1a-1* mutant plants, as indicated. Note that the *cdt1a-1* mutant is lethal and therefore the CDT1a-CFP protein seems to be fully functional.

**Figure S3.**
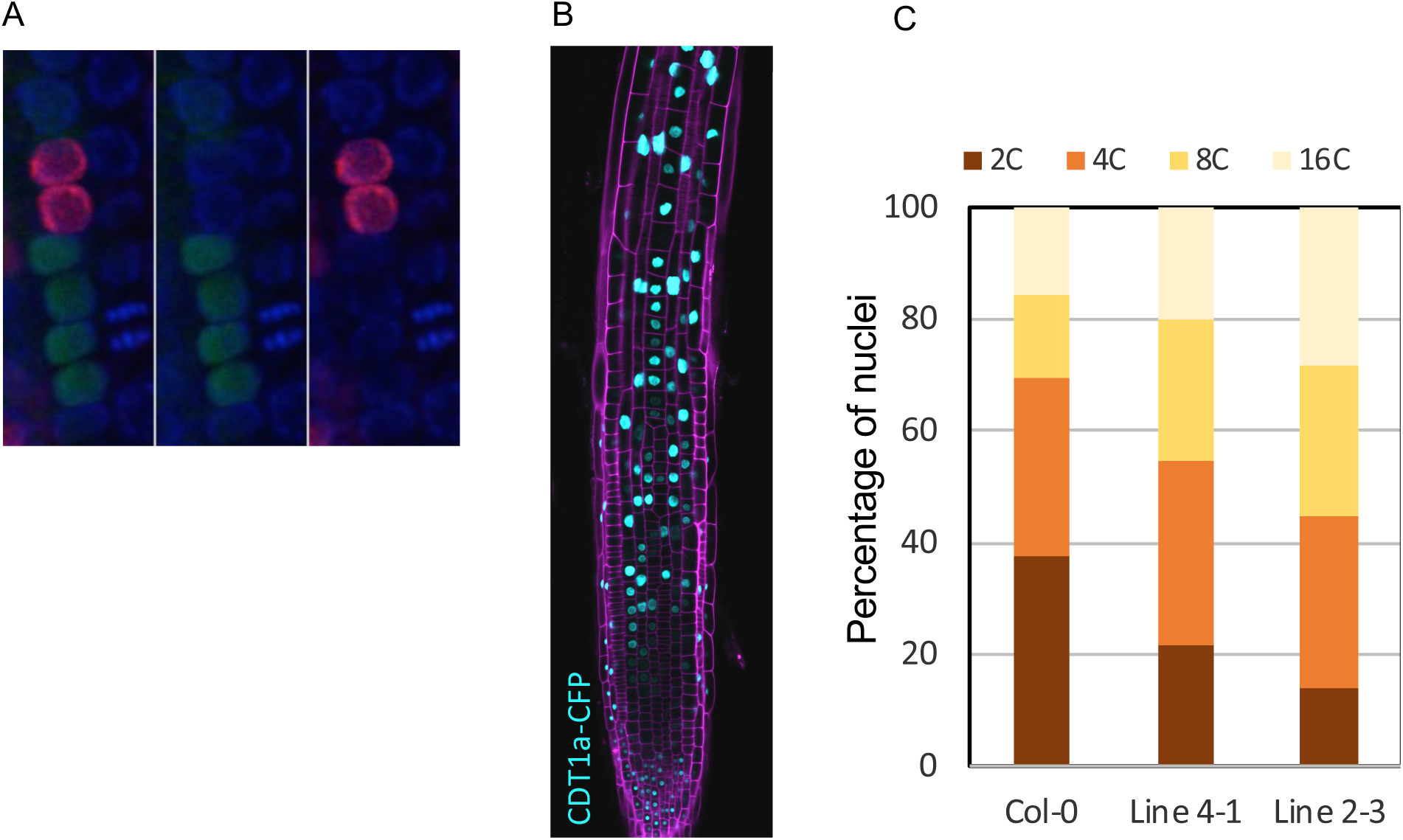
Constitutive expression of CDT1a-CFP. (A) Constitutively expressed CDT1a is degraded right after the G1/S transition, as it is the case when expressed from its own promoter. Note that EdU labeling and CFP signals do not colocalize. (B) The expression domain of constitutively expressed CDT1a-CFP in the root apex is extended to the differentiated part of the root. (C) The nuclear ploidy profile is different from control plants since there is a reduction of 2C nuclei and an increase in 16C nuclei, likely the result of ectopic expression of CDT1a-CFP in the transition/differentiation zone, where cells normally endoreplicate, and not in proliferating cells.

## Additional files

**Supplementary Table 1.** Primers used in this study

### Supplementary Movies

**Movie supplement 1.** Loading and degradation of CDT1a-CFP during G1.

**Movie supplement 2.** Dynamics of CDT1a-mRFP and FBL17-GFP.

---

**Supplemental Table 1.**
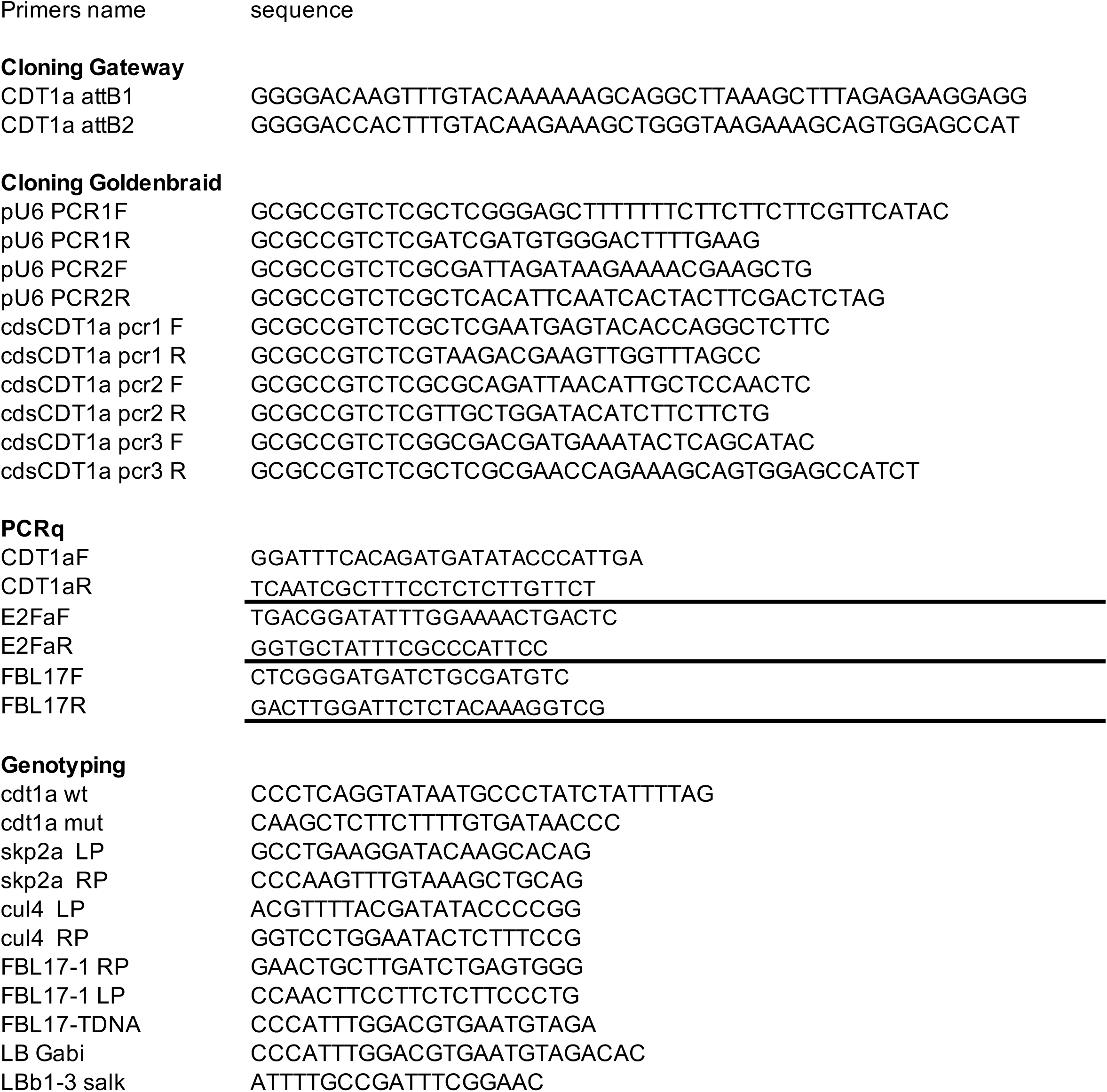
Description of primers used in this study.

